# Proton Conductivity of Glycosaminoglycans

**DOI:** 10.1101/388686

**Authors:** John Selberg, Manping Jia, Marco Rolandi

## Abstract

Proton (H^+^) conductivity is important in many natural phenomena including oxidative phosphorylation in mitochondria and archea, uncoupling membrane potentials by the antibiotic Gramicidin, and proton actuated bioluminescence in dinoflagellate. In all of these phenomena, the conduction of H^+^ occurs along chains of hydrogen bonds between water and hydrophilic residues. These chains of hydrogen bonds are also present in many hydrated biopolymers and macromolecule including collagen, keratin, chitosan, and various proteins such as reflectin. All of these materials are also proton conductors. Recently, our group has discovered that the jelly found in the Ampullae of Lorenzini-shark’s electrosensing organs- is the highest naturally occurring proton conducting substance. The jelly has a complex composition, but we attributed the conductivity to the glycosaminoglycan keratan sulfate (KS). Here, we have measured the proton conductivity of hydrated keratan sulfate using PdH_x_ contacts to be 0.50 ± 0.11 mS cm ^-1^- consistent to that of Ampullae of Lorenzini jelly, 2 ± 1 mS cm ^-1^. Proton conductivity, albeit with lower values, is also shared by other glycosaminoglycans with similar chemical structures including dermatan sulfate, chondroitin sulfate A, heparan sulfate, and hyaluronic acid. This observation confirms the structure property relationship between proton conductivity and the chemical structure of biopolymers.

## Introduction

Proton (H^+^) conductivity is important in many natural phenomena(1) including oxidative phosphorylation in mitochondria and archea(2-4), uncoupling membrane potentials by the antibiotic Gramicidin(5), and proton actuated bioluminescence in dinoflagellate(6). In all of these phenomena, the conduction of H^+^ occurs along chains of hydrogen bonds between water and hydrophilic residues. These chains are often referred to as proton wires(3). This conduction follows the Grotthus mechanism in which a hydrogen bond is exchanged with a covalent bond contributing to the effective transfer of an H^+^ from a molecule to its next-door neighbor(7). Following this mechanism, proton conductivity in hydrated biopolymers and macromolecules is widespread including collagen(8), keratin(9), chitosan(10), melanin(11), peptides(12), and various proteins such as bovine serum albumin(13) and reflectin(14, 15). In addition to the ability to support proton wires, typically these materials include an acid or a base group that serve as H^+^ or OH^-^ dopants and provide charge carriers for proton conductivity (16-18). Following this trend, for example, the synthetic polymer Nafion, with a high proton conductivity of 78 mS cm^-1^, contains very strong acid groups that donate H^+^ to the water of hydration for proton conduction (19). Our group has recently demonstrated that the jelly contained in the ampullae of Lorenzini, the electrosensing organ of sharks and skates, is the highest naturally occurring proton conductor(20). We postulated that keratin sulfate (KS), a glycosaminoglycan (GAG) was the material responsible for proton conductivity due to its similarity to chitosan and Nafion, and the ability form proton wires when hydrated. Here, we have measured the proton conductivity of KS derived from bovine cornea (21, 22) and other GAGs using Pd based proton conducting devices (10).

## Materials and methods

### Materials

Glycosaminoglycan samples were received from the Linhardt laboratory at Rensselaer University and stored dry at -15C. Including, 70% pure CSA isolated from bovine trachea (average MW: 20kDa), HA sodium salt from streptococcus zooepidemicus (average MW: 100kDa), DS from porcine intestinal mucosa from porcine intestinal mucosa (average MW: 30kDa), HS (porcine intestinal mucosa (average MW: 14.8kDa), and KS isolated from the bovine cornea (average MW: 14.3kDa).

### Device Fabrication

Two-terminal measurements were performed on Si substrates with a 100-nm SiO_2_ layer. Conventional photolithography was used to pattern Pd contacts. Pd contacts were 500 um wide and 100 nm thick separated by different channel lengths, L_SD_= 5, 10, 20, 50, 100, 200, 500 um. We performed both two terminal device measurements and transmission line measurements (TLM) to reduce the influence of contact resistance on the conductivity (11).

### Deposition of Glycosaminoglycans

All samples were dissolved in DI water at a ratio of 0.15 – 0.2 mg ul^-1^ and drop casted onto the devices. The samples were then dried with dry nitrogen gas flow.

### Proton Conductivity Measurements

Direct current–resistance measurements were performed using a Keithley 4200 source-meter and a two-contact probe station arrangement on devices. The devices were enclosed in an environmental chamber at room temperature in an atmosphere of nitrogen or hydrogen with controlled relative humidity (RH) at 75%, 90%, 90% with 5% hydrogen, and 90% with 5% deuterium gas to form PdHx or PdDx contacts. A one-hour incubation period was carried out after switching between humidity and gas compositions. During the measurement, the Pd/PdHx electrodes were contacted with tungsten probes. When we applied a source-drain potential difference, VSD, the PdHx source injected protons (H&^#x002B;^) into drain through the samples, inducing measurable electrical current in the circuit.

## Results and Discussion

GAGs are long, linear, hydrophilic biopolymers composed of repeating of disaccharide units with many acidic groups that may support the presence of proton wires (Fig. 1b), including hyaluronic acid (HA), heparan sulfate (HS), chondroitin sulfate A (CSA), dermatan sulfate (DS), and keratan sulfate (KS). Additionally, GAGs have important biological functions in regulating hydration and water homeostasis of tissues, which is derived from their ability to absorb very large amounts of water at high humidity(23).

**Figure 1.**
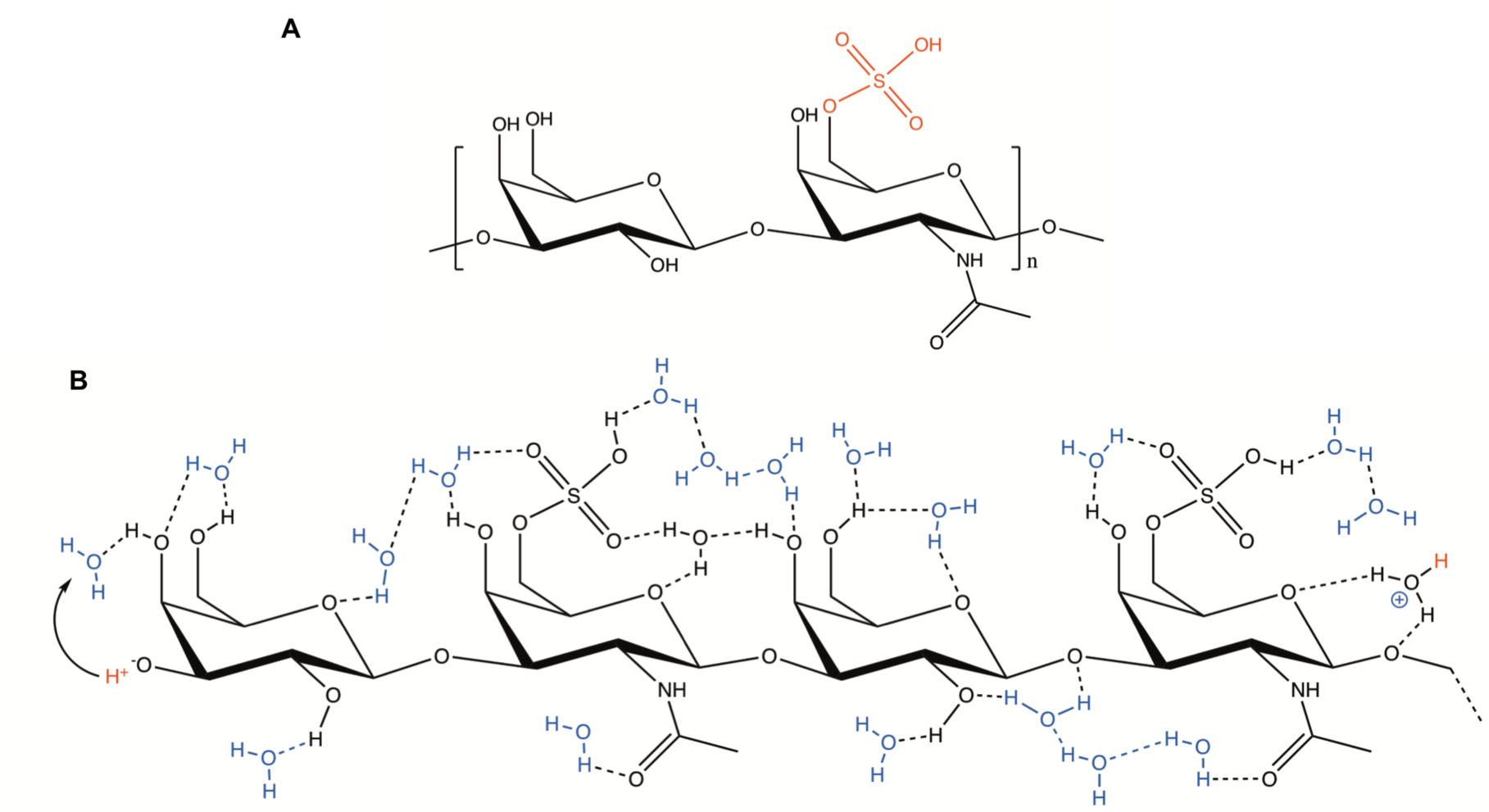
The keratan sulfate. (A) Chemical structure of keratan sulfate. (B) An illustration of a three-monomer segment of keratan sulfate. Possible intra- and inter-molecular hydrogen bonds as well as the hydrogen bonds between the water of hydration and the polar parts of the molecule form a continuous network comprised by hydrogen-bond chains. The sulfate group interacts with the hydrogen-bond network and forms an H_3_O^+^ (hydronium) ion.

### Proton Conductivity Measurements

Palladium (Pd) devices are useful for studying proton transport in materials due to the nature of Pd to reversibly form palladium hydride (PdH_x_)(24-27). Several mechanisms for the formation of PdH_x_ are known (equations 1-4).

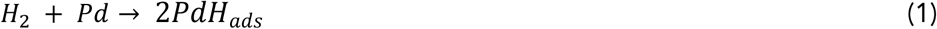

Equation one describes the adsorption and splitting of H_2_ molecules into two adsorbed H on the Pd metal surface without electron transfer in a reaction described by Tafel kinetics.

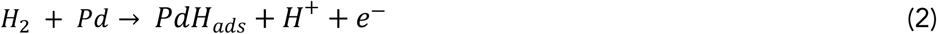

Equation two is the Heyrovsky reaction in which a H_2_ is split into an adsorbed H atom and a H^+^, e^-^ pair at the Pd surface, this e^-^ is transferred into the metal.

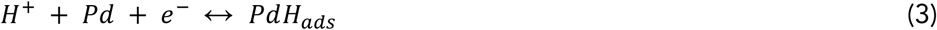

The Volmer reaction in equation 3 describes a third mechanism, which involves an electron transfer to a H^+^ near the Pd surface allowing it to adsorb as PdH_ads_. Once PdH_ads_ is formed on the metal surface, H can diffuse into the subsurface bulk forming PdH_x_ (eq. 4) . (10, 28, 29).

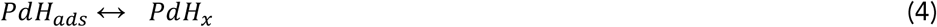

Pd devices were designed such that PdH_x_ formation occurs spontaneously by equation 1 in a 5% H_2_ atmosphere on two Pd contacts. These Pd/PdH_x_ contacts are separated by a channel consisting of a GAG film (Figure 2a, 2b). A voltage V_SD_ between the Pd/PdH_x_ contacts induces a current of H^+^ to exit one Pd contact, travel through the GAG channel, and enter the second Pd contact according to equation 3. In this manner, one e^-^ travels through the circuit and is recorded as Id for each H^+^ that is conducted through the GAG channel.

**Figure 2.**
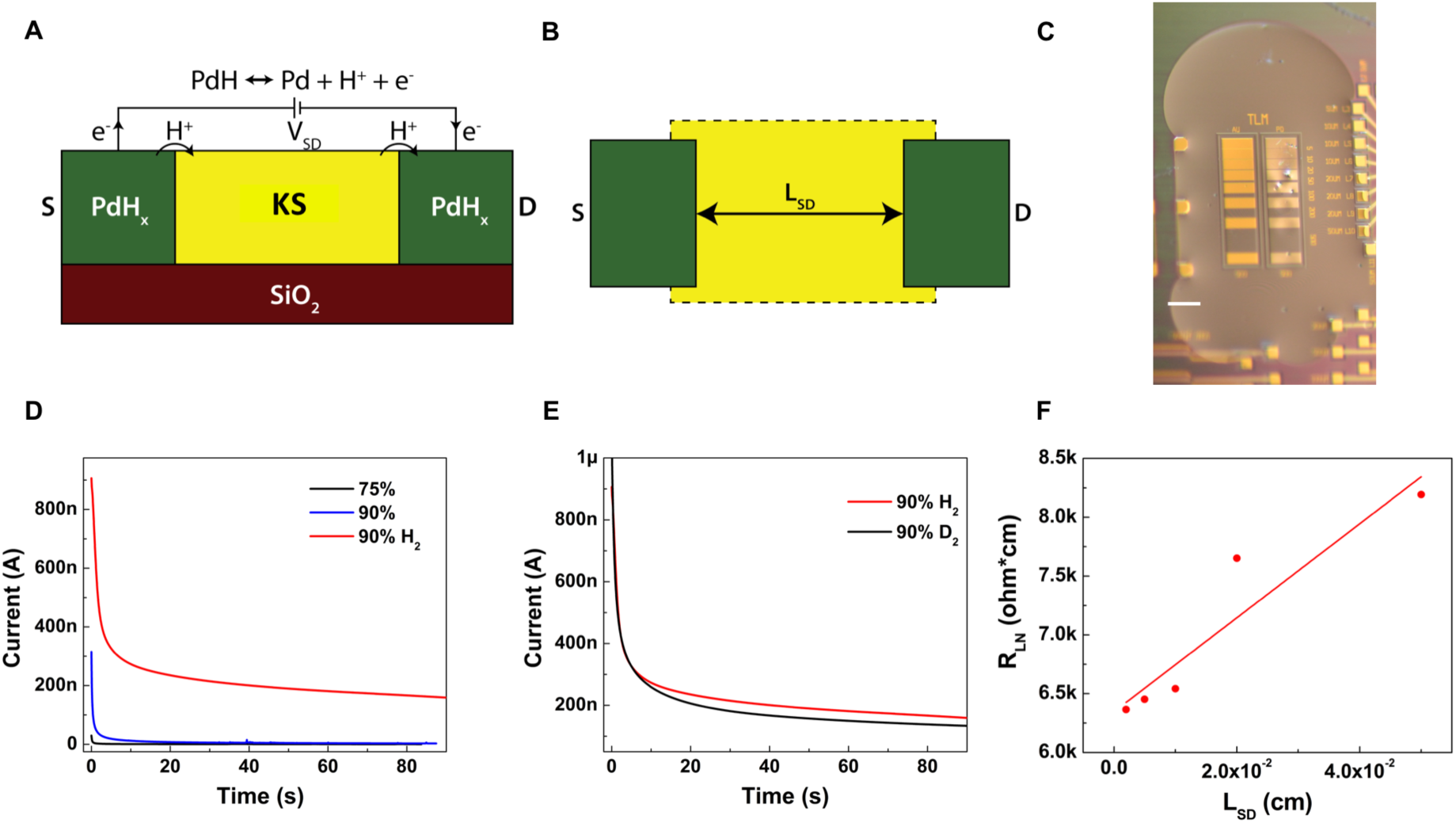
Proton conduction measurement of keratan sulfate. **A)** Palladium hydride(PdH_x_) electrode behavior. Under a V_SD_, PdH_x_ source split into Pd, H^+^, and e^−^. Protons are injected into the KS, whereas electrons travel through external circuitry and are measured. **B)** TLM geometry. Varying the distance between source and drain (L_SD_) distinguishes between the fixed PdH_x_ – KS interface contact resistance and the varying bulk resistance. **C)** Optical image of TLM geometry with hydrated KS on the surface. Scale bar, 0.5 mm. **D)** Transient response to a 1V bias in KS at 75%, 90%, 90% H_2_ RH, in which the current under 90% with hydrogen is much higher than that under 90% RH without hydrogen. **E)** Deuterium current (black) at 90% D2 humidity is lower than proton current (red). **F)** RLN as a function of L_SD_, A linear fit gives a bulk material proton conductivity of 0.50 ± 0.11 mS cm^-1^.

### Materials characteristics

After deposited directly onto the transmission line measurement (TLM) (Figure 2c) device surface without further processing, the KS film is thick, viscous, and optically transparent. After one hour of incubating at 50%RH, the KS film dries to a non-homogenous film. The film rehydrates fully after incubating at 90%RH for one hour and appears as wet as when it was drop-cast form solution (Fig. 2C). This high water content of KS films is a result of sulfate groups functionalizing either or both of the galactose and N-acetyl glucosamine sugars which make up the repeating disaccharide unit of the GAG. Furthermore, these hydrophilic sulfate groups may induce the organization of water into hydrogen bond chains much like the sulfate group in the commercial proton exchange polymer Nafion. This organization of water allows proton hopping along the chain according to the Grotthuss mechanism(30). Considering the other members of GAGs family, DS, HS, CSA, HA also contain an abundance of repeating acidic groups which may stabilize proton wires, as shown in Table S1, we performed a screen for their proton conductivity characteristics.

### DC electrical measurements with PdH_x_ proton-conducting contacts

With V_SD_ =1V on the Pd devices, we measured the drain current (I_D_) of KS, as shown in Fig 2D. At RH = 75% in pure nitrogen, I_D_ ~ 0.5 nA is small (black). With the RH increasing to 90% in pure nitrogen, the increase in I_D_ was negligible (red). However, after the addition of H_2_ to the gas mixture, I_D_ increased more than 300 times to 155 nA (green). The two orders of magnitude increase after the inclusion of hydrogen to the atmosphere, which promotes the formation of PdH_x_, which is proton conducting, indicates a strong component of proton conductivity. We also performed the same measurement with other materials in GAGs family and observed similar trends (Fig. S1). All GAGs displayed a high increased current upon a 90%RH with 5%H_2_ atmosphere compared to a 90%RH N_2_ atmosphere, indicating that protons predominately contribute to the conductivity of GAGs materials at high relative humidity.

### Kinetic isotope effect

To further verify that KS conductivity predominantly arises from protons, we investigated the kinetic isotope effect. Measurements were repeated while hydrating the sample with deuterium oxide (D_2_O) instead of water and exposing the sample to deuterium gas rather than H_2_. Like protons, deuterium ions (D^+^) can transport along proton wires and hydrated materials, albeit with a lower mobility and an associated lower current because of the higher molecular weight(31). The kinetic isotope effect in KS is evident as a drop in the conductivity when deuterium replaces hydrogen as the atom being transported (Fig 2E). Here, we observe a 15% drop in current when deuterium replaces hydrogen. The kinetic isotope effect observed with KS is relatively small. However, a similar small kinetic isotope effect was observed for the proton conduction of bovine serum albumin(13). The other members in GAGs family display a larger kinetic isotope effect, the current drop is nearly 50% (Fig. S2).

### Transmission line measurement

TLM devices are designed with different lengths between the Pd source and the drain contacts to eliminate the effect of contact resistance in the measurements of the proton conductivity (Fig 2B) (20). We applied V_SD_ = 1 V on devices with L_SD_ ranging from 5 to 500 um, measured I_D_, and calculated the resistance of each device, R_L_. In this geometry, RL increases linearly with L_SD_, but the contact resistance, R_C_, at the source–KS and drain–KS interface is constant. Considering that different devices contained KS with different thicknesses, we multiplied RL by the sample thickness to get the normalized resistance. The slope of the plot of R_LN_ as a function of L_SD_ is proportional to the resistivity of KS, and the intercept on the R_LN_ axis for L_SD_ = 0 is R_CN_ (Fig 2F). Here, we obtain σ = 0.50 ± 0.11 mS cm^-1^, which is only one order of magnitude lower than the proton conductivity of Nafion σ = 58.3 ± 2.5 mS cm^-1^(20) measured in the same geometry (Fig S3). The proton conductivity of the Nafion control sample (58.3 ± 2.5 mS cm^-1^) measured in a TLM geometry is extremely close to the reported value of 78 mS cm^-1^.(19) Therefore, we conclude that σ = 0.50 ± 0.11 mS cm^-1^ measured in this way is a good indicator of the proton conductivity of KS. Table 1 shows the proton conductivity of Nafion and known biopolymers, and KS performs well among them.

**Table 1.**
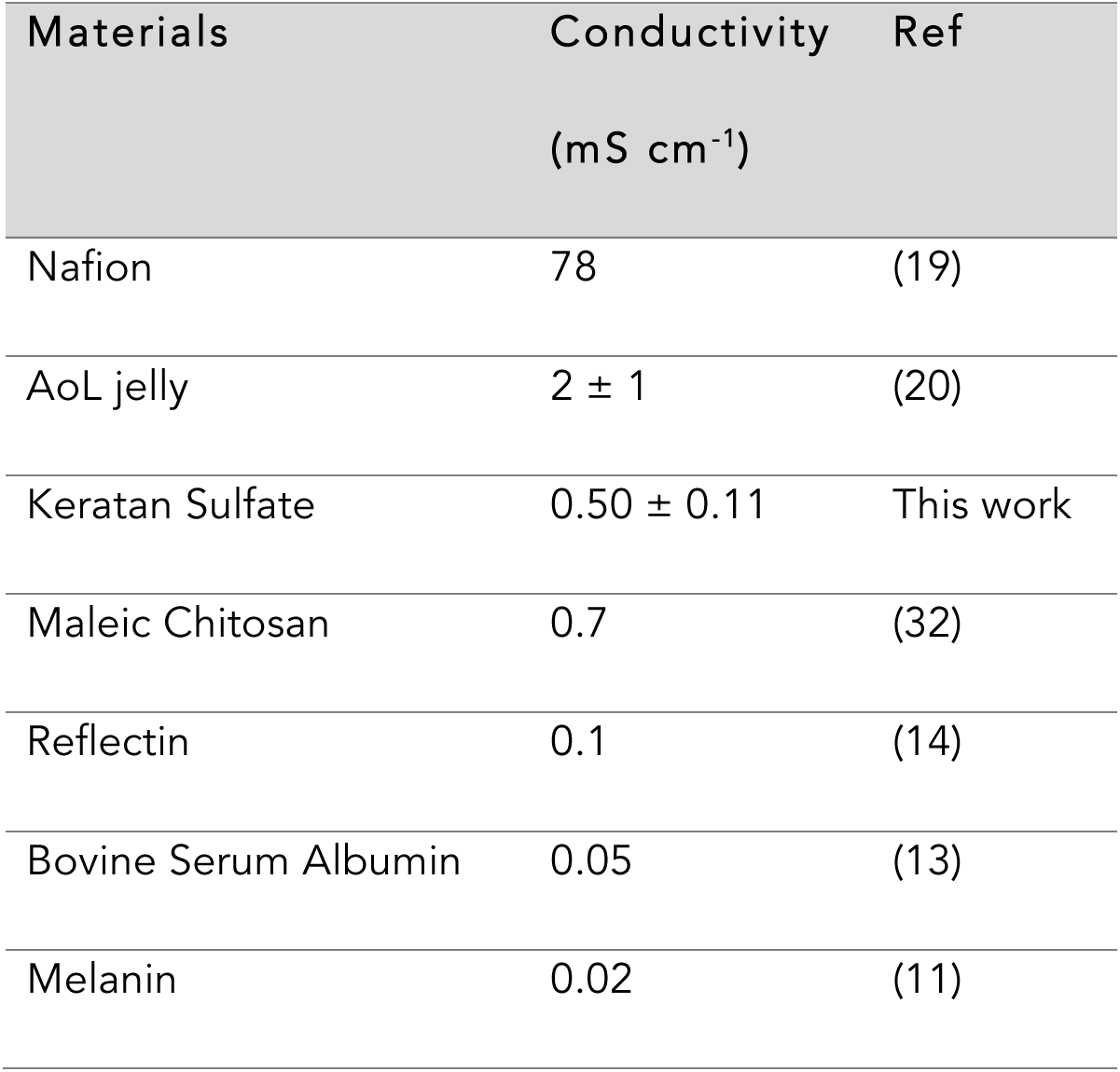
Room-temperature proton conductivities of Nafion and known biopolymers.

We have also measured the proton conductivities of other GAGs reported in table S1. Hyularonic acid has the highest conductivity 0.28 ± 0.06 mS cm^-1^. However, some of the other GAGs materials, such as dermatan sulfate, did not form a homogeneous film and it was not possible to measure the conductivity using the TLM geometry. The conductivity reported with the two terminal geometry also contains contact resistance and therefore it is lower as expected. All of these GAG have acidic group that provide H^+^ for proton conductivity. Within experimental error, we did not observe any variation in conductivity with variation in pK_a_. It is difficult to relate the concentration of H^+^ in these hydrated states because pK_a_ is determined in infinite dilution. If we assume that with the water of hydration is at neutral pH, we can expect all of the acidic groups on the polymers to be ionized independent of pK_a_.

## Conclusions

Inspired by the high conductivity in the jelly of the ampullae of Lorenzini, we measured the proton conductivity of Keratan Sulfate and other glycoaminoglicans with similar chemical structures. Using TLM devices at room temperature, we measured the proton conductivities of 0.50 ± 0.11 mS cm^-1^ at 90% RH H_2_ for KS, which is comparable to that of ampullae of Lorenzini jelly (2 ± 1 mS cm^-1^). This result confirms our assumption that the high proton conductivity of the ampullae of Lorenzini jelly arises mainly from keratan sulfate. We have also measured the proton conductivity of other GAGs including heparan sulfate, dermatan sufate, chondroitin sulfate A and hyaluronic acid. Their conductivity is lower, but comparable with keratan sulfate suggesting that proton conductivity is a common property of GAGs with acidic groups when hydrated.

---

**Figure SI 1.**
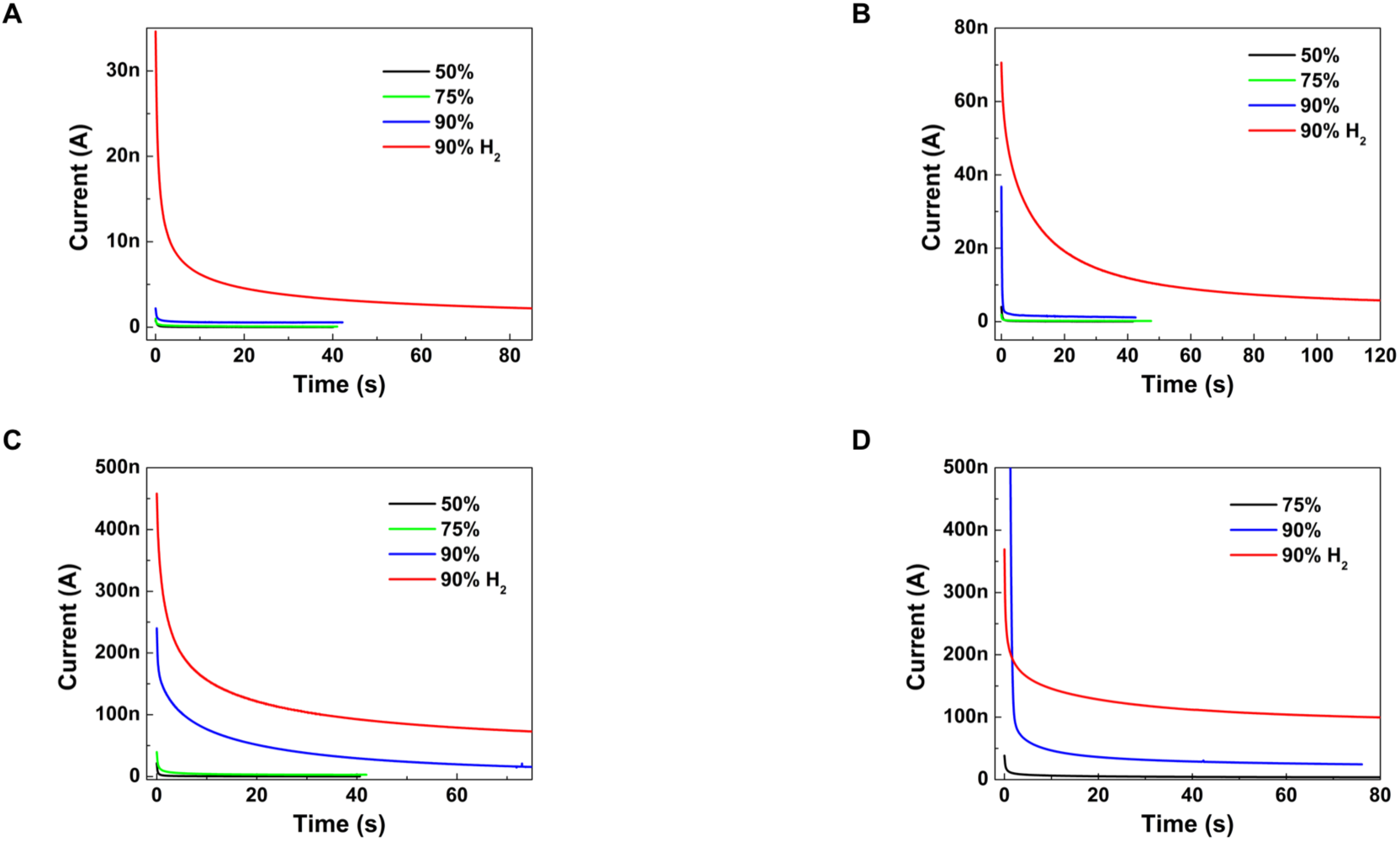
Current under different RH of GAG family. **(A)** hyaluronic acid, **(B)** heparan sulfate, **(C)** chondroitin sulfate A and **(D)** dermatan sulfate. The current under 90%RH with hydrogen is much higher than 90%RH without hydrogen.

**Figure SI 2.**
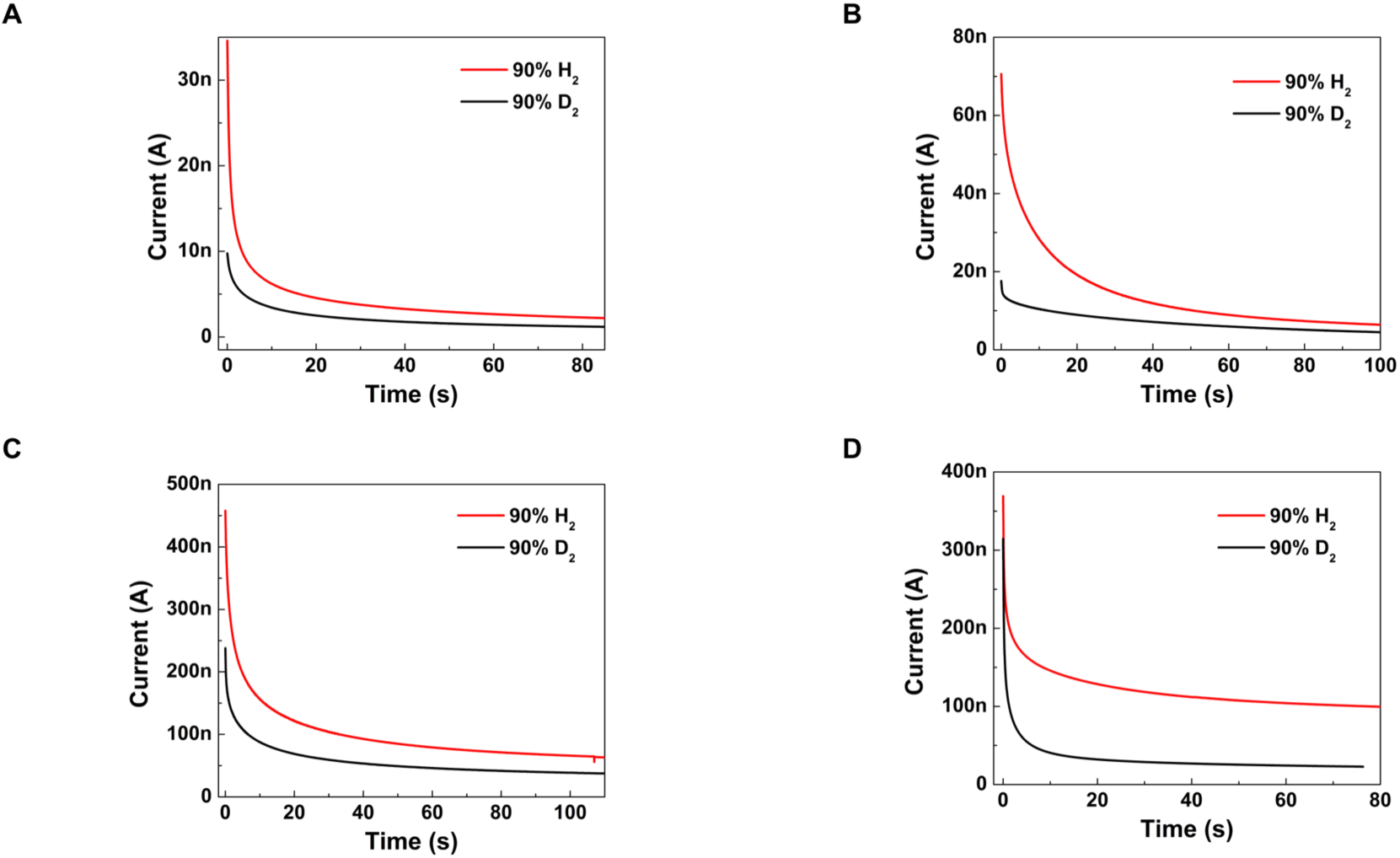
Kinetic isotope effect in members of GAG family. **(A)** hyaluronic acid, **(B)** heparan sulfate, **(C)** chondroitin sulfate A and **(D)** dermatan sulfate. Current measured in a 5% deuterium (black) atmosphere at 90%RH vs a 5% proton atmosphere at 90%RH (red).

**Figure SI 3.**
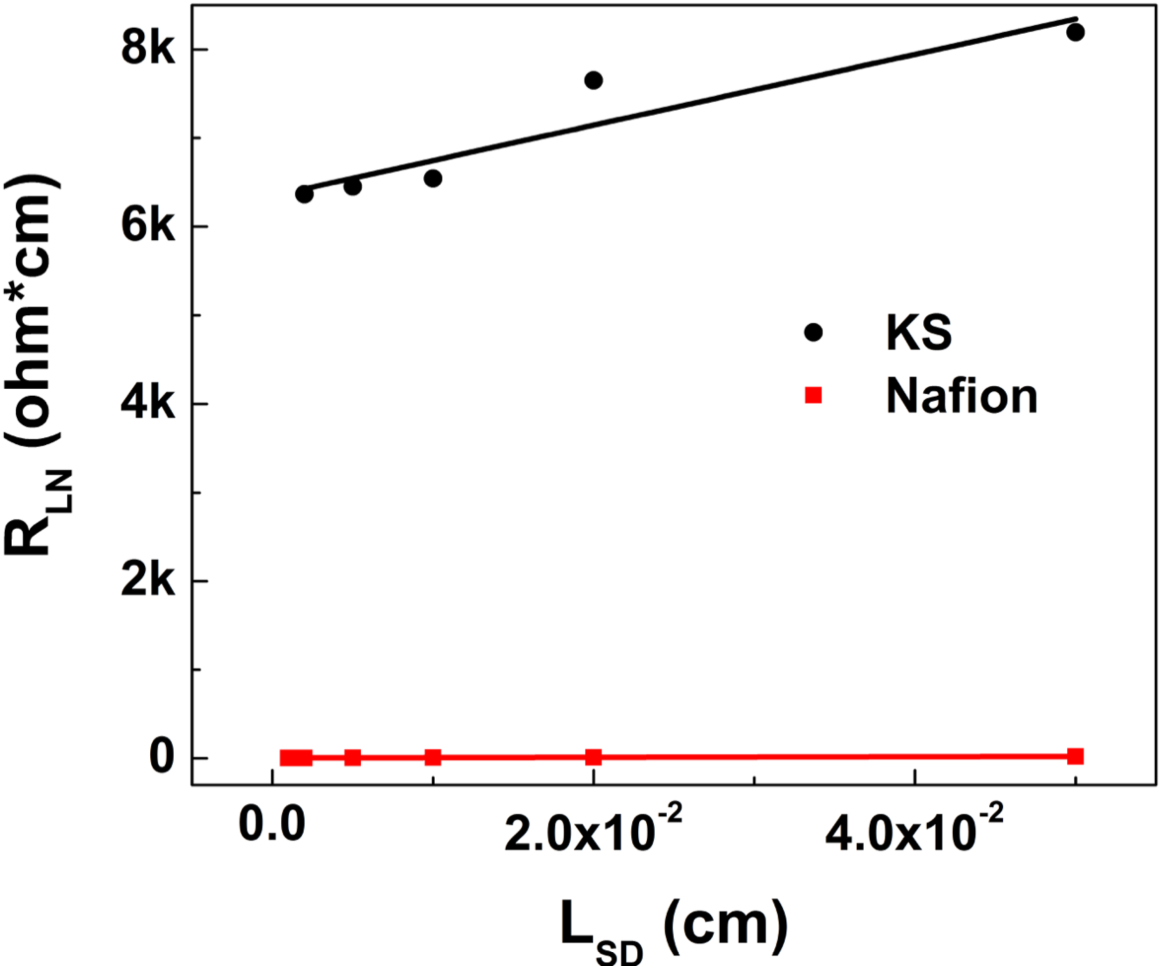
Control experiments on Nafion. Conductivity of Nafion measured with this TLM device is 58.3 ± 2.5 mS cm^-1^. It’s slightly lower than the literature value of 78 mS cm^-1^, which is attributed to sample preparation. The reported literature value is after immersion in heated sulfuric acid, while the sample here was simply drop-cast from solution.

**Table S1.**
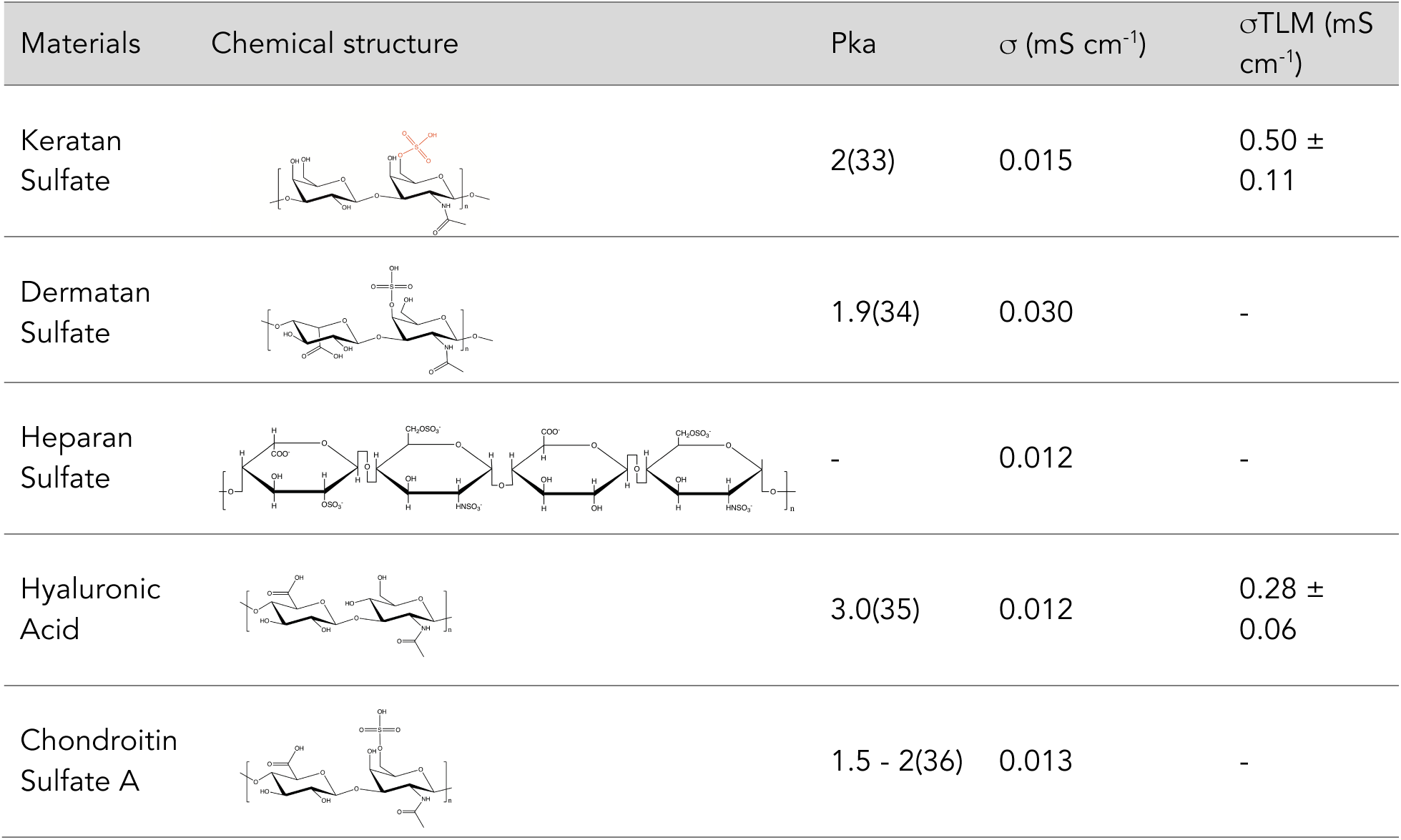
Glycosaminoglycan chemical structures, Pka and conductivity (σ) estimated with TLM devices

| Materials | Chemical structure | Pka | $\sigma$ (mS cm<sup>-1</sup>) | $\sigma_{\text{TLM}}$ (mS cm<sup>-1</sup>) |
| --- | --- | --- | --- | --- |
| Keratan Sulfate | | 2(33) | 0.015 | $0.50 \pm 0.11$ |
| Dermatan Sulfate |  | 1.9(34) | 0.030 | - |
| Heparan Sulfate |  | - | 0.012 | - |
| Hyaluronic Acid |  | 3.0(35) | 0.012 | 0.28 ± 0.06 |
| Chondroitin Sulfate A |  | 1.5 - 2(36) | 0.013 | - |

